# A broad and potent H1-specific human monoclonal antibody produced in plants prevents influenza virus infection and transmission in guinea pigs

**DOI:** 10.1101/2020.01.13.902841

**Authors:** Jun-Gyu Park, Chengjin Ye, Michael S. Piepenbrink, Aitor Nogales, Haifeng Wang, Michael Shuen, Ashley J. Meyers, Luis Martinez-Sobrido, James J. Kobie

## Abstract

Although seasonal influenza vaccines block most predominant influenza types and subtypes, humans still remain vulnerable to waves of seasonal and new potential pandemic influenza viruses for which no immunity may exist because of viral antigenic drift and/or shift, respectively. Previously, we have described a human monoclonal antibody (hMAb), KPF1, which was produced in human embryonic kidney 293T cells (KPF1-HEK) with broad and potent neutralizing activity against H1N1 influenza A viruses (IAV) *in vitro*, and prophylactic and therapeutic activities *in vivo*. In this study, we produced hMAb KPF1 in tobacco plants (KPF1-Antx) and demonstrate how the plant-produced KPF1-Antx hMAb possesses similar biological activity compared with the mammalian produced KPF1-HEK hMAb. KPF1-Antx hMAb shows broad binding to recombinant HA proteins and H1N1 IAV, including A/California/04/2009 (pH1N1) *in vitro*, that are comparable to those observed with KPF1-HEK hMAb. Importantly, prophylactic administration of KPF1-Antx hMAb to guinea pigs prevented pH1N1 infection and transmission in both prophylactic and therapeutic experiments, substantiating its clinical potential to prevent and treat H1N1 infections. Collectively, this study demonstrates, for the first time, that plant-produced influenza hMAbs have similar *in vitro* and *in vivo* biological properties to those produced in mammalian cells. Because of the many advantages of plant-produced hMAbs, such as rapid batch production, low cost, and the absence of mammalian cell products, they represent an alternative strategy for the production of immunotherapeutics for the treatment of influenza viral infections, including emerging seasonal and/or pandemic strains.

## 1. Introduction

Influenza viruses are members of the *Orthomyxoviridae* family and responsible for severe respiratory disease in humans [1]. Influenza viruses cause both seasonal epidemics and occasional pandemics of great consequence when novel viruses are introduced into the human population [2]. Although Food and Drug Administration (FDA)-licensed inactivated and live-attenuated influenza vaccines have been used for over seventy years, yearly seasonal influenza viral infections are still responsible for an estimated of 3 to 5 million severe cases of illness and approximately 250,000 to 500,000 annual deaths [1, 3–6]. Influenza virus public health concerns are further aggravated by their ability to efficiently transmit and by the limited available antivirals [7–11]. Influenza viruses are divided into types A, B, C, and D [12–14]. Influenza A viruses (IAV) are further classified into different subtypes based on the antigenic surface glycoproteins hemagglutinin (HA; eighteen subtypes) and neuraminidase (NA; eleven subtypes) [1, 15–18]. Among them, HA subtypes are classified into group 1 (H1, H2, H5, H6, H8, H9, H11, H12, H13, and H16-18) and group 2 (H3, H4, H7, H10, H14, and H15) subtypes [15, 16, 19]. Currently, H1N1 and H3N2 IAV and influenza B viruses (IBV) are circulating in the human population and responsible for seasonal influenza [1, 3]. Among the different influenza viruses, H1N1 IAVs are the most important subtype and have been responsible for causing pandemics with enormous impact on both human health and economy, as illustrated by the 1918 and 2009 pandemics [1].

Currently, FDA vaccines and antivirals are available for the prevention and treatment, respectively, of influenza viral infections in humans. Influenza vaccines contain viral antigens representing the prevalent H3N2 and H1N1 IAVs as well as one (trivalent) or two (quadrivalent) lineages (Victoria or Yamagata) of IBV currently circulating in humans [20–22] and are provided as inactivated or live-attenuated forms. However, although influenza vaccines are able to induce immunity, they have several limitations. These include their lack of effectiveness against drifted seasonal viruses, as recently illustrated with the pandemic H1N1 virus in 2009, the lag time (∼2 weeks) to establish an effective immune responses after exposure, and their limited, if any, protection against shifted pandemic viruses. Moreover, yearly formulations are required to protect against constantly changing IAV and/or IBV. In terms of antivirals, three major classes of drugs are FDA-approved for the treatment of influenza infections: NA inhibitors (oseltamivir, zanamivir and peramivir), matrix protein 2 (M2) inhibitors (amantadine and rimantadine), and the polymerase acid (PA) endonuclease inhibitor (Baloxavir marboxil or Xofluza) [1, 23]. However, influenza antiviral drugs have several limitations, including the lack of antiviral activity of M2 inhibitors against IBV, the emergence of drug resistant variants [24–26], and a limited antiviral effect due to rapid metabolism and elimination of the inhibitor [27–30]. Thus, there is an urgent need to find alternative approaches for the prevention and treatment of influenza infections in humans.

Since 1990, plants have been considered as a potential biofactory for production of biologics, including monoclonal antibodies (MAbs) [31, 32]. MAbs produced in genetically engineered plants (or plantibodies) [33] have been shown to have similar biological activity compared to mammalian cell-produced MAbs [34, 35]. Because large quantities of MAbs are generally required for passive immunization *in vivo*, plantibodies represent an excellent option for the production of MAbs for passive immunization, including large-scale manufacturing, low cost and the absence of animal components [33, 35–37]. Moreover, plantibodies overcome some of the concerns associated with animal-derived therapeutic MAbs obtained from serum or plasma, including intermediate reactions, pyrogenicity, potential contamination with other zoonotic pathogens and/or toxins, and serum sickness [33, 35–38]. For these reasons, plantibodies have been proposed for the treatment of several bacterial (e.g. Salmonella, Streptococcus, Porphyromonas) and viral (e.g. Ebola virus, EBOV; Hepatitis B virus, HBV; Human immunodeficiency virus, HIV; Porcine epidemic diarrhea virus, PEDV; Rabies virus, RABV; West Nile virus, WNV) infections; or toxins (e.g. Ricin and Shiga toxin) [36, 39–47]. However, to date, plantibodies have not been used for the treatment of influenza viral infections.

In a recent study, we described that a human monoclonal antibody (hMAb) KPF1 produced in mammalian human embryonic kidney (HEK293T) cells (KPF1-HEK) showed broadly cross-reactive activity against H1N1 IAV *in vitro* [48]. This is because KPF1-HEK hMAb recognizes a highly conserved and novel epitope in the HA1 globular head region of H1 IAV with high affinity. Importantly, KPF1-HEK hMAb showed broad and potent neutralizing activity *in vitro* and prophylactic and therapeutic activity in mice against influenza H1N1 IAV [48]. In this study, we demonstrate that a plant-produced KPF1 hMAb (KPF1-Antx) exhibits broad cross-reactivity and potent neutralizing *in vitro* and *in vivo* activity against H1N1 IAV. Importantly, KPF1-Antx hMAb has prophylactic and therapeutic activity and could prevent viral transmission in the well-established guinea pig model of influenza virus infection. Notably, the biological properties of KPF1-Antx hMAb are similar to those of the HEK293T-produced KPF1 hMAb [19]. Altogether, our results demonstrate, for the first time, the feasibility of using plantibodies for the treatment of influenza viral infections in humans.

## 2. Materials and Methods

### 2.1. Cells and Viruses

Madin-Darby Canine Kidney (MDCK; ATCC CCL-34) and human embryonic kidney (HEK293T; ATCC CRL-3216) cells were maintained in Dulbecco’s modified Eagle’s medium (DMEM; Mediatech, Inc.) supplemented with 5% fetal bovine serum (FBS) and 1% PSG (100 unit/ml penicillin, 100 μg/ml streptomycin, and 2 mM L-glutamine) at 37°C in a 5% CO_2_ atmosphere [19]. Wild-type (WT) influenza viruses A/Brisbane/59/2007 H1N1 (Brisbane/H1N1), A/ California/04/2009 H1N1 (pH1N1), A/New Caledonia/20/1999 H1N1 (NC/H1N1), A/Puerto Rico/8/1934 H1N1 (PR8/H1N1), A/Wisconsin/629-D02473/2009 H1N1 (WI/H1N1), A/Georgia/F32551/2009 H1N1 (GA/H1N1), and A/Brisbane/10/2007 H3N2 (Brisbane/H3N2) were propagated in MDCK cells as previously described [48–50].

### 2.2. Production of KPF1 in HEK293T cells

HEK293T cells were grown at 37°C to approximately 80% confluence in 10 cm tissue culture dishes using DMEM with 5% Hyclone Fetal Clone II (GE Healthcare Lifesciences, Logan, UT) and 1X antibiotic and antimycotic (Gibco, Grand Island, NY). Expression plasmids containing the heavy and light chain sequences for KPF1 were transfected into HEK293T cells using jetPRIME transfection reagent (PolyPlus, New York, NY) as previously described [19]. Media was harvested and replenished three times over 8 days. IgG was purified from culture supernatant using Magne Protein A beads (Promega, Madison, WI) and the elution buffer exchanged with PBS using Amicon Ultra centrifugal filters (Millipore-Sigma, Cork, Ireland).

### 2.3. Production of KPF1 in plants

Plant expression vectors were assembled using standard recombinant DNA technology, as previously described [51]. The KPF1-Antx and isotype control antibody expression vectors, which included heavy chain (HC) and light chain (LC) genes, were co-expressed with an oligosaccharyltransferase from Leishmania major (LmSTT3D) known to enhance N-glycan occupancy of recombinant proteins, using a similar strategy to that previously described [52]. In addition, a third expression vector was utilized to enhance recombinant protein expression by transiently silencing Argonaute1 (AGO1) and Argonaute4 (AGO4) proteins to minimize post-transcriptional gene silencing (PTGS) [Patent WO 2019/023806 A1].

KPF1-Antx was produced as outlined in Figure 1A. Briefly, expression vectors were transformed into *Agrobacterium tumefaciens* strain EHA105 and infiltrated at an OD_600_ of 0.2 into *Nicotiana benthamiana* plant line KDFX, developed by PlantForm (unpublished) for knock-down of the plant-specific β1,2-xylosyltransferase and α1,3-fucosyltransferase [53]. Plant foliage was harvested 7 days post-infiltration and total soluble protein extracted. Antibodies were purified using MabSelect Protein A followed by Capto Q according to manufacturer protocols (GE Healthcare, Chicago, IL). Purified antibodies were concentrated and formulated to ≥25 mg/mL in PBS.

**Figure 1.**
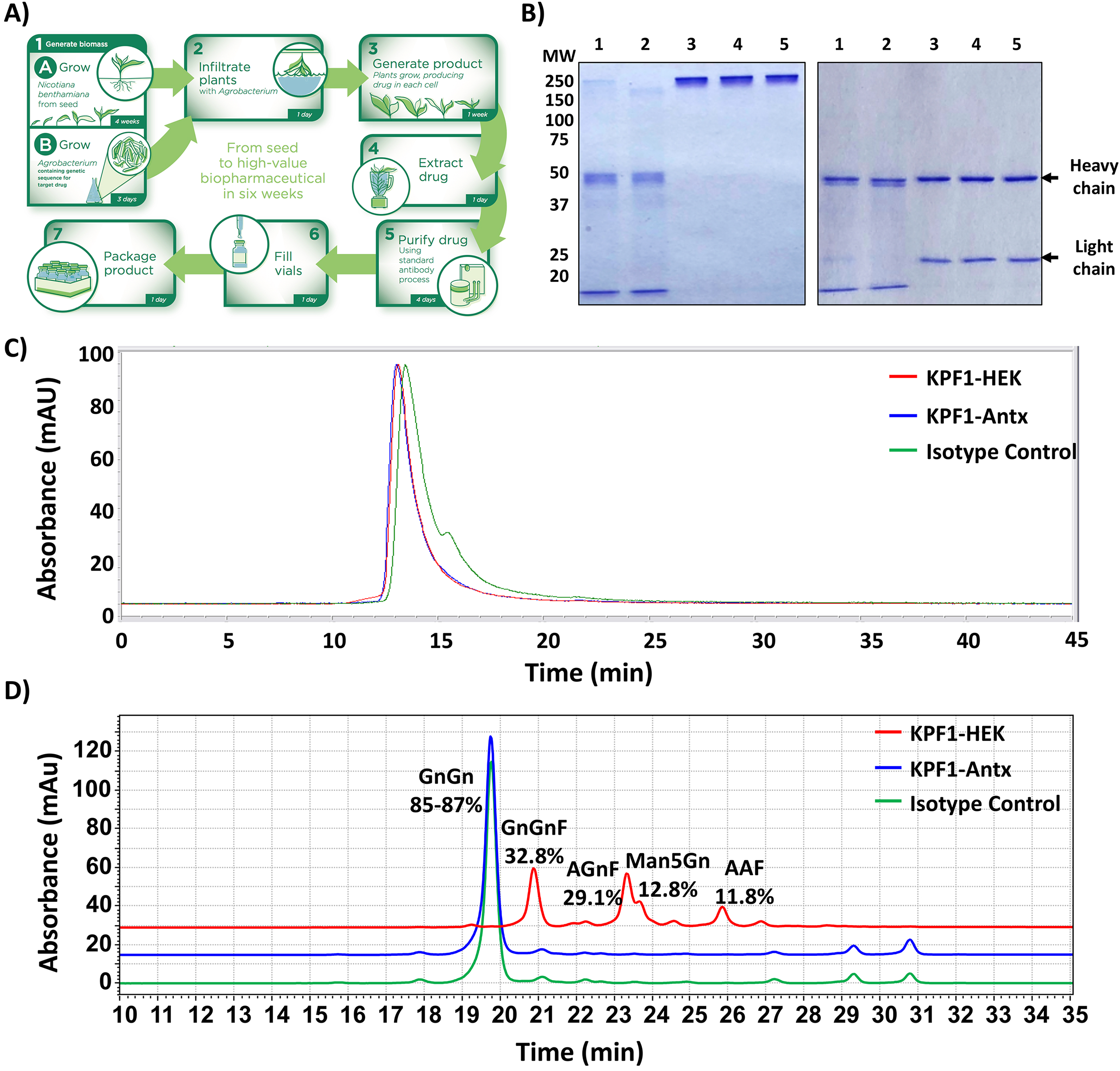
Production and characterization of the KPF1-Antx hMAb. **A) Schematic representation of KPF1-Antx hMAb production using the *vivo*XPRESS^®^ platform.** Four-week-old plants were vacuum-infiltrated with a solution containing *Agrobacterium tumefaciens*. The Agrobacteria contained binary vectors for the co-expression of KPF1 heavy and light chain genes, STT3D and a system for enhancing protein expression. The infiltrated plants were returned to the greenhouse for seven days, after which, all foliage was harvested and processed to purify KPF1 MAb using a standard antibody process. **B) KPF1-Antx hMAb purification process and analysis:** Samples from the KPF1-Antx hMAb purification process: Protein A load (1), protein A flow through (2), protein A elution (3), CaptoQ FT (4), and final purified KPF1-Antx hMAb (5) were analyzed under non-reducing (left) and reducing (right) conditions using standard SDS-PAGE electrophoresis and Coomassie blue staining. Mobility of the molecular marker (MW) in kDa is shown on the left. **C) Size exclusion high performance liquid chromatography:** Purified hMAbs (10 µg) were injected into a TSKgel G3000SWXL column equipped with a TSKgel guard column SWXL (Tosoh Biosciences) using an Agilent 1100 Series HPLC. Agilent LC/MSD ChemStation Edition software was used for integration of data. **D) N-glycan analysis of KPF1-Antx and KPF1-HEK hMAbs, and isotype control:** Glycans were prepared using the GlykoPrep® Rapid N-Glycan Preparation kit (PROzyme) and separated by Hydrophilic-Interaction Liquid Chromatography (HILIC) using a TSKgel Amide-80 column (Tosoh Bioscience). Glycan species were identified by relative retention time and quantified using autointegration of each glycan species peak.

### 2.4. Coomassie blue staining and Western blot

A total amount of 2.5, 1.25, 0.625 and 0.313 μg of KPF1-HEK or KPF1-Antx hMAbs were mixed with loading buffer and phosphate-buffered saline (PBS) to a final volume of 20 μl. After separated by 12% SDS-PAGE, the gel was stained with Coomassie blue staining solution (0.05% Coomassie brilliant blue R-250, 45% Methanol and 7% Acetic acid) overnight at room temperature. Alternatively, the gel was transferred to a nitrocellulose membrane. After blocking with 5% bovine serum albumin (BSA) in PBS containing 0.1% Tween 20 (PBST) at room temperature for 1 h, the membrane was incubated with horseradish peroxidase-conjugated goat anti-human IgG secondary antibody (Jackson ImmunoResearch Laboratories, PA, USA) at room temperature for another 1 h. The blot was developed with ECL detection reagent (Thermo Fisher, CA, USA) in the ChemiDoc MP Imaging System (BioRad, PA, USA).

### 2.5. Enzyme-linked immunosorbent assay (ELISA)

Binding of KPF1-HEK or KPF1-Antx hMAbs to H1N1 or H3N2 HA proteins was performed using standard ELISA, as previously described [19]. Briefly, ELISA plates (Nunc Maxisorp, Thermo Fisher Scientific, Grand Island, NY) were coated overnight with 1 µg/ml of the indicated recombinant HA proteins obtained from Biodefense and Emerging Infectious Research Resources Repository (BEI Resources, Manassas, VA) and incubated with serially 10-fold diluted HEK-or Antx-produced KPF1 hMAbs in PBS (starting concentration of 10 µg/ml). Binding was detected with HRP-conjugated anti-human IgG (Jackson ImmunoResearch, PA, USA). In selected ELISAs, increasing concentrations of urea (ranging from 0 to 8 M) were added and the plates incubated for 15 min at room temperature prior to detection with anti-IgG-HRP to evaluate avidity.

### 2.6. Immunofluorescence assay (IFA)

MDCK cells (96 well plate format, 5 x 10^4^ cells/well, triplicates) were inoculated with Brisbane/H1N1, pH1N1, NC/H1N1, PR8/H1N1, WI/H1N1, GA/H1N1, and Brisbane/H3N2 at multiplicity of infection (MOI) of 10, or mock infected. At 8 h post-infection (p.i.), cells were fixed with 10% neutral buffered formalin (NBF; Thermo Fisher Scientific, CA, USA) for 1 h and then stained with 1 μg/ml of KPF1-HEK or KPF1-Antx hMAbs, followed by secondary Alexa Fluor 488-conjugated goat anti-human IgG antibody (Life Technologies, Carlsbad, CA, USA). DAPI (4’,6-diamidino-2-phenylindole, Thermo Fisher Scientific, CA, USA) was used to stain cell nuclei. Fluorescent signal images were acquired under an inverted fluorescent microscope (Olympus, Japan) and analyzed by ImageJ software to measure intensity of the staining (NIH, Bethesda, MD, USA) [54].

### 2.7. Microneutralization assays (MNA)

Virus MNA were performed as previously described [19]. Briefly, KPF1-HEK or KPF1-Antx hMAbs, or IgG1 isotype controls, were serially 2-fold diluted in PBS in 96-well plates (starting concentration of 200 µg/ml). One hundred plaque forming units (PFU) of each virus (Brisbane/H1N1, pH1N1, NC/H1N1, WI/H1N1, GA/H1N1, and Brisbane/H3N2) were then added to the hMAb dilutions and incubated for 1 h at room temperature. MDCK cells (96-well plate format, 5 × 10^4^ cells/well, quadruplicates) were then infected with the hMAb-virus mixture for 1 h at room temperature. After viral adsorption, cells were maintained in p.i. medium, with 1 µg/ml of *N*-tosyl-_L_-phenylalanine chloromethyl ketone (TPCK)-treated trypsin (Sigma), and incubated at 37°C. Virus neutralization was determined by crystal violet staining at 72 h p.i. The neutralization titer 50 (NT_50_) was determined by a sigmoidal dose response curve (GraphPad Prism, v7.0) [19]. Mock-infected cells and viruses in the absence of the KPF1 hMAbs were used as internal control.

### 2.8. Hemagglutination inhibition (HAI) assays

HAI assays were performed to determine HA-neutralizing capability of KPF1-Antx and KPF1-HEK hMAbs, as previously described [19]. Briefly, KPF1-Antx or KPF1-HEK hMAbs were serially 2-fold diluted (staring concentration of 200 µg/ml) in 96-well V-bottom plates and mixed 1:1 with 4 hemagglutinating units (HAU) of Brisbane/H1N1, pH1N1, NC/H1N1, PR8/H1N1, WI/H1N1, GA/H1N1, or Brisbane/H3N2 for 60 min at room temperature. HAI titers were determined by adding 0.5% turkey red blood cells (RBCs) to the virus-hMAb mixtures for 30 min on ice. HAI titers were defined as the minimum amount of hMAb that completely inhibited hemagglutination.

### 2.9. In vivo experiments

Four-week-old female Hartley guinea pigs were purchased from Charles River Laboratory and maintained in the animal care facility at University of Rochester under specific pathogen-free conditions. All animal protocols were approved by the University of Rochester Committee of Animal Resources and complied with the recommendations in the Guide for the Care and Use of Laboratory Animals of the National Research Council [55]. For viral infections, guinea pigs were anesthetized intraperitoneally (i.p.) with Ketamine (30 mg/kg) and Xylazine (5 mg/kg) and inoculated intranasally (i.n.) with 10^3^ PFU of pH1N1 in 100 μl of PBS. After viral infection, animals were monitored daily for morbidity (body weight loss) and mortality (survival) (data not shown). To determine the prophylactic efficacy of KPF1-Antx hMAb, guinea pigs in two groups (N=3/group) were weighed and administered (i.p.) 20 mg/kg of KPF1-Antx hMAb, or isotype control hMAb (isotype-Antx) and kept in separated cages. After six h, guinea pigs were infected (i.n.) with 10^3^ PFU of pH1N1. One day post-infection (d p.i.), sentinel guinea pigs (N=3/group) were introduced into the cages of infected guinea pig and monitored for seven days. Viral replication in nasal washes at 2, 4, 6, and 8 d p.i. was determined by immunofocus assay (fluorescent focus-forming units, FFU/ml) using an anti-NP MAb (HB-65) and a FITC-conjugated anti-mouse secondary Ab (Dako). Geometric mean titers and data representation were performed using GraphPad Prism, v7.0. For therapeutic efficacy, guinea pigs in two groups (N=3/group) were infected (i.n.) with 10^3^ PFU pH1N1 and kept in separated cages, followed by administration of KPF1-Antx or Isotype-Antx hMAbs at 1 d p.i. One day after administration of hMAbs, sentinel guinea pigs (N=3/group) were introduced into the cages of infected guinea pig and monitored for seven days. Viral replication in nasal washes collected at 2, 3, 5, 7, and 9 d p.i. was determined by immunofocus assays as described above. At the completion of the study, guinea pigs were humanely euthanized by administration of a lethal dose of avertin and exsanguination, and lungs were collected for gross observation. Macroscopic pathology scoring was evaluated using ImageJ software to determine the percent of the total surface area of the lung (dorsal and ventral view) affected by consolidation, congestion, and atelectasis as previously described [54, 56, 57].

### 2.10. Statistical analysis

The one-tailed unpaired Student’s t-test was used to evaluate significant differences. Data were expressed as the mean ± standard deviation (SD) using Microsoft Excel software. Values were considered statistically significant when * *p* < 0.05, ** *p* < 0.01, or no significance (n.s.). All data were analyzed using Prism software version 8.00 (GraphPad Software, CA, USA).

## 3. Results

### 3.1. Production of the human monoclonal antibody KPF1 in tobacco plants

KPF1-Antx was produced in four-week old *N. benthamiana* plants as outlined in Figure 1A. Just one week after infiltration with transgene carrying *Agrobacterium tumefaciens*, IgG was purified from foliage using Protein A followed by Capto Q to remove impurities including endotoxin. KFP1-Antx was expressed at an average of 650 mg/kg of biomass (n=3, data not shown) and the overall recovery was 68% with an endotoxin level of 0.4 endotoxin units (EU)/mg. Antibody recovery and quality were monitored throughout the purification process using standard SDS-PAGE and Coomassie Blue staining (Figure 1B). IgG can be observed in the Protein A load in Figure 1B in addition to host cell proteins such as RuBisCO, which can account for up to 50% of total soluble proteins in leaves [58]. The final KPF1-Antx product is reduced to two independent bands representing the heavy and light chains (50 and 25 kDa, respectively) with no impurities detected. KPF1-Antx and KPF1-HEK were compared using size exclusion HPLC analysis (Figure 1C). Area under the curve analysis indicated that KPF1-Antx contained 96.3% monomeric IgG and 3.4% low molecular weight (MW) forms; whereas, KPF1-HEK contained 94.5% monomeric IgG, 3.9% low MW forms and 1.6% high MW forms. These results indicate a greater purity for the plant-derived KPF1-Antx, which included a polishing step (Capto Q). In addition, N-glycosylation profiles were compared using GlykoPrep® Rapid N-Glycan Preparation kit (PROzyme, Hayward, CA) and separation by Hydrophilic-Interaction Liquid Chromatography (HILIC) using a TSKgel Amide-80 column (Figure 1D). KPF1-Antx and the isotype control N-glycan profiles were highly similar, with 85-87% biantennary N-acetylglucosamine (GnGn). Contrary, KPF1-HEK N-glycan profile contained a mixture of N-glycans typically observed on mammalian glycoproteins, including antibodies (32.8% GnGnF, 29.1% AGnF, 12.8% Man5Gn and 11.8% AAF).

### 3.2. Reactivity of KPF1-Antx and KPF1-HEK hMAbs in vitro

We initially characterized the KPF1 hMAbs generated from either HEK293T cells (KPF1-HEK) or tobacco plants (KPF1-Antx) *in vitro* (Figure 2). Serially 2-fold diluted (2.5 µg to 0.313 μg) KPF1-HEK and KPF1-Antx hMAbs showed similar characteristics by SDS-PAGE (Figure 2A) and Western blot (Figure 2B), regardless of mammalian or plant production. A comprehensive binding assessment of KPF1-Antx and KPF1-HEK hMAbs to various IAV HA proteins, including Brisbane/H1N1, pH1N1, NC/H1N1, PR8/H1N1, A/Christchurch/16/2011 H1N1 (ChCh/H1N1), A/St. Petersburg27/2011 H1N1 (St. Petersburg/H1N1), and Brisbane/H3N2 was performed (Figure 2C). As expected, KPF1-Antx hMAb bound only H1 HAs including the more recent ChCh/H1N1 and St. Petersburg/H1N1, and binding of KPF1-Antx hMAb to different IAV H1s was similar to that of KPF1-HEK hMAb (Figure 2C). To evaluate the stability of the binding, two different concentrations (0.1 and 1 µg/ml) of KPF1-Antx and KPF1-HEK hMAbs were treated with increasing concentrations of urea (Figure 2D). Both KPF1-Antx and KPF1-HEK hMAbs maintained similar binding affinity in 4 M urea, and substantially diminished in 8 M urea (Figure 2D). These results indicate similar biological properties of the mammalian- and plant-produced KPF1 hMAbs *in vitro*.

**Figure 2.**
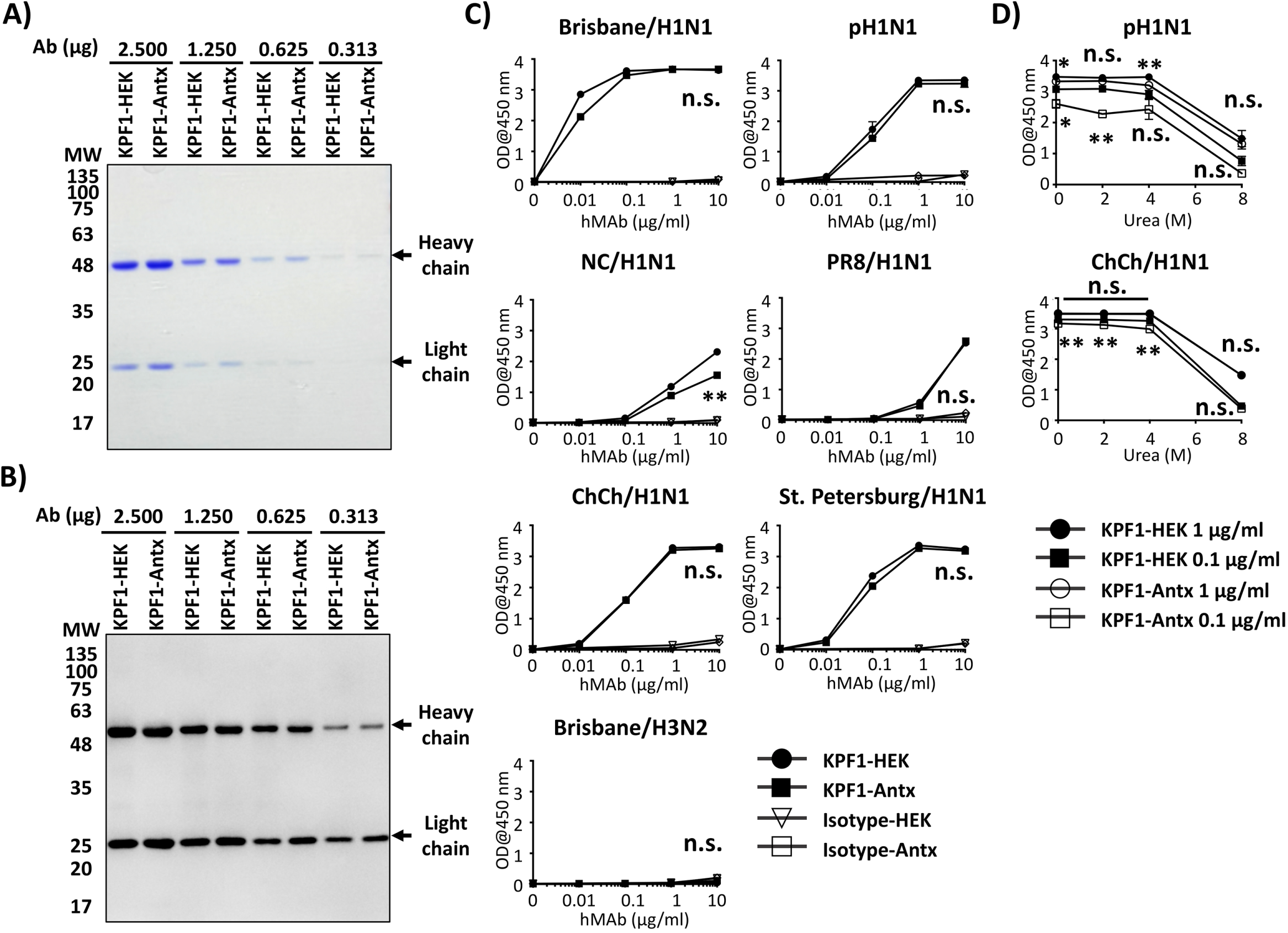
*In vitro* characterization of KPF1-Antx hMAb. **A-B) Purity and mobility of the KPF1-HEK and KPF1-Antx hMAbs:** Equal amount of KPF1 hMAbs (2.500, 1.250, 0.625 and 0.313 μg) generated from either HEK293T cells (KPF1-HEK) or tobacco plant (KPF1-Antx) were separated by SDS-PAGE and evaluated by Coomassie blue staining (**A**) or Western blot using an anti-human IgG HRP-conjugated Ab (**B**). **C) Recombinant HA binding of KPF1-HEK and the KPF1-Antx hMAbs:** Binding to 1 μg /ml of indicated HA proteins was determined for serially diluted hMAbs by ELISA. n.s.: No significance. **D) Avidity:** Binding of hMAbs in the presence of increasing concentrations of urea was determined by ELISA. The statistical analysis between KPF1-HEK and KPF1-Antx (markers above the lines; between 1 μg/ml of KPF1-HEK and KPF1-Antx, marker below the lines: between 0.1 µg/ml of KPF1-HEK and KPF1-Antx), * *p* < 0.05, ** *p* < 0.01, or no significance (n.s.).

### 3.3. Cross-reactivity of KPF1-Antx hMAb to H1N1 IAV

To characterize the ability of the KPF1-HEK and KPF1-Antx hMAbs to recognize native IAV HA proteins, MDCK cells were infected (MOI 10) with Brisbane/H1N1, pH1N1, NC/H1N1, PR8 /H1N1, WI/H1N1, GA/H1N1, or Brisbane/H3N2 and binding was evaluated and quantified by IFA (Figure 3). Both KPF1-Antx and KPF1-HEK hMAbs recognized all H1N1 IAVs, but not Brisbane/H3N2 or mock-infected cells (Figure 3A). Quantifying the KPF1-HEK and KPF1-Antx hMAbs binding reactivity to infected MDCK cells revealed that KPF1-Antx hMAb showed similar levels of recognition and binding to H1N1-infected MDCK cells as that observed with KPF1-HEK hMAb (Figure 3B). This result indicates that KPF1-Antx hMAb possesses similar broad cross-reactivity and ability to recognize HA from H1N1 IAV-infected MDCK cells as those observed with the mammalian HEK-produced KPF1 hMAb.

**Figure 3.**
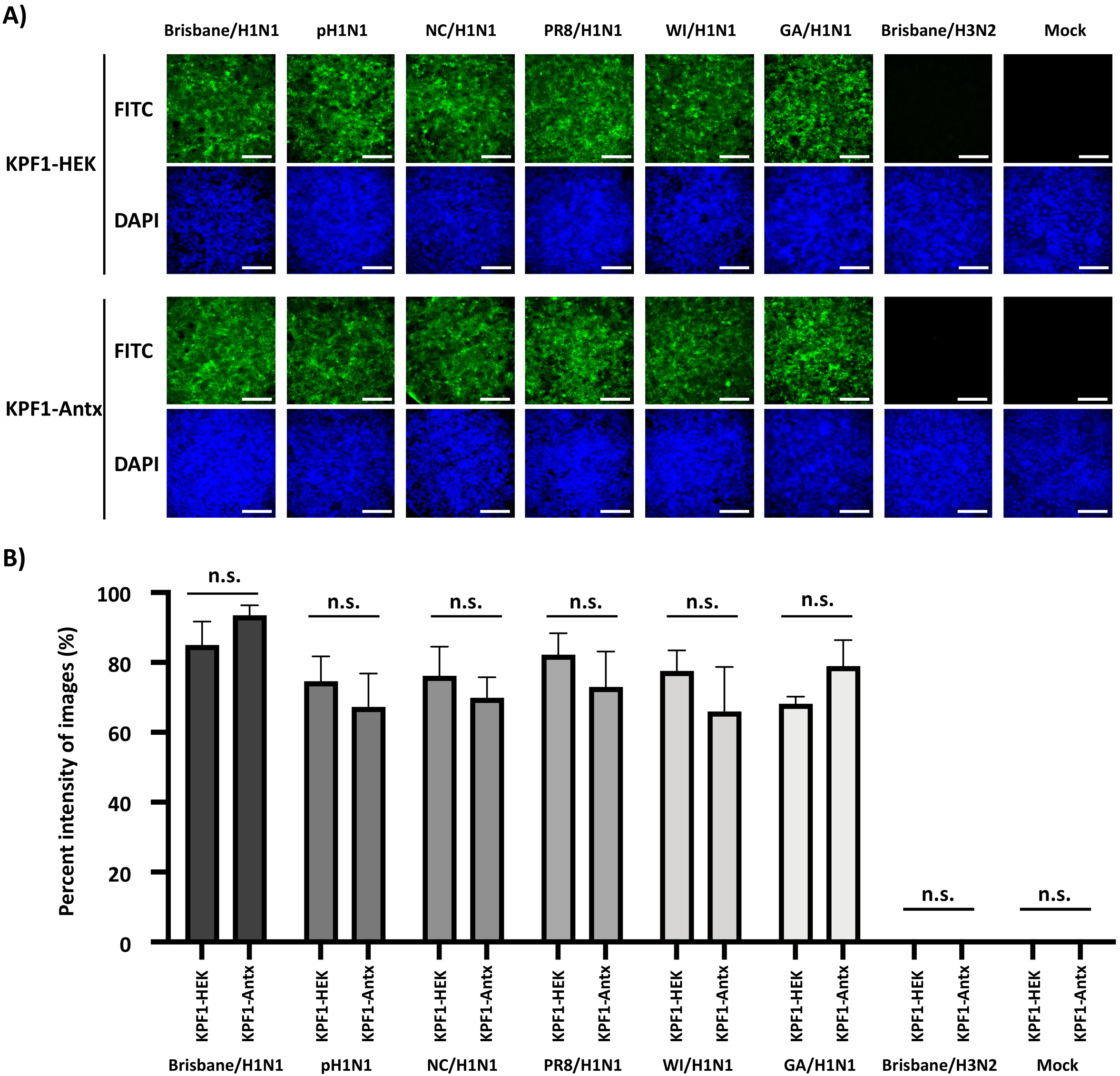
Cross-reactivity of KPF1-HEK and KPF1-Antx hMAbs against IAV H1N1 strains. **A) IFA:** MDCK cells were infected (MOI 10) with Brisbane/H1N1, pH1N1, NC/H1N1, PR8/H1N1, WI/H1N1, or GA/H1N1; and Brisbane/ H3N2 influenza A virus; or mock infected. At 12 h p.i., cells were fixed and the cross-reactivity of the KPF1-HEK (top) or KPF1-Antx (bottom) hMAbs were evaluated by IFA. DAPI was used for nuclear staining. Scale bar = 100 µm. **B) Cross-reactivity intensity**: Images in panel A were analyzed for the intensity of fluorescence using ImageJ. n.s.: No significance.

### 3.4. Broad neutralization and hemagglutination inhibition activity by KPF1-Antx hMAb

We next evaluated the ability of KPF1-Antx hMAb to neutralize a broad range of H1N1 IAV, similar to previously described with KPF1-HEK hMAb [19]. Both KPF1-Antx and KPF1-HEK hMAbs showed similar cross-reactivity, as assessed by IFA, against different H1N1 viruses (Figure 3A). Importantly, quantification of the IFA results indicates that the ability of KPF1-Antx hMAb to recognize the IAV H1 in infected cells is not statistically different than that of KPF1-HEK hMAb (Figure 3B). Likewise, KPF1-Antx hMAb is able to similarly neutralize Brisbane/H1N1, pH1N1, NC/H1N1, WI/H1N1, and GA/H1N1 strains (NT_50_ KPF1-Antx = 0.195 to 0.780 μg/ml; NT_50_ KPF1-HEK = 0.195 to 1.106 μg/ml), while neutralization of PR8/H1N1 was slightly less efficient (NT_50_ KPF1-Antx = 12.5 μg/ml; NT_50_ KPF1-HEK = 25.0 μg/ml) (Table 1). As expected, KPF1 hMAbs were not able to neutralize Brisbane/H3N2 IAV, even at the highest concentration (200 μg/ml). Both KPF1-Antx and KPF1-HEK hMAbs also showed similar HAI activity against H1N1 IAV and were potent for Brisbane/H1N1, pH1N1, WI/H1N1, and GA/H1N1(HAI = 0.271 to 1.106 μg/ml) but less efficient for NC/H1N1 and PR8/H1N1 (HAI = 8.714 to 17.425 μg/ml) strains (Table 1).

**Table 1.** MNA and HAI assays with KPF1-HEK and KPF1-Antx hMAbs.

| Viruses | NT<sub>50</sub> (µg/ml) MNA |  | HAI (µg/ml) |  |
| --- | --- | --- | --- | --- |
|  | KPF1-HEK | KPF1-Antx | KPF1-HEK | KPF1-Antx |
| **Brisbane/H1N1** | 0.708 | 0.780 | 0.552 | 0.552 |
| **pH1N1** | 0.427 | 0.427 | 1.106 | 0.552 |
| **NC/H1N1** | 1.106 | 0.715 | 8.714 | 17.425 |
| **PR8/H1N1** | 25.0 | 12.5 | 17.425 | 17.425 |
| **WI/H1N1** | 0.195 | 0.195 | 0.271 | 0.271 |
| **GA/H1N1** | 0.353 | 0.221 | 0.271 | 0.271 |
| **Brisbane/H3N2** | >200 | >200 | ND | ND |

### 3.5. Prophylactic activity of KPF1-Antx hMAb in vivo

To evaluate the protective efficacy of KPF1-Antx hMAb, Hartley guinea pigs (N=3/group) received 20 mg/kg of either KPF1-Antx or isotype-Antx control hMAbs 6 h prior to infection with 10^3^ PFU of pH1N1 (Figure 4A). Sentinel guinea pigs (N=3/group) were put in contact with the infected guinea pigs at 1 d p.i. to assess viral transmission. In the KPF1-Antx hMAb treated group, only one infected guinea pig and its sentinel guinea pig showed restricted viral infection and shedding, respectively, in the nasal washes (Figures 4B and C). Contrarily, all infected guinea pigs treated with the isotype-Antx hMAb control showed higher levels of viral replication (Figure 4B). Also, we observed viral shedding that resulted in efficient transmission to sentinel guinea pigs in all animals treated with the isotype-Antx hMAb (Figure 4C). Importantly, gross pathology supported that KPF1-Antx hMAb has prophylactic activity (Figures 4D-4F). Both, infected and sentinel guinea pigs in the KPF1-Antx hMAb treated group showed mild or no multifocal consolidation, congestion and atelectasis in middle and caudal lobes (Figures 4D and E, respectively), while the isotype-Antx hMAb control treated group infected (Figure 4D) and sentinel (Figure 4E) guinea pigs showed more severe pathology. Distribution of pathologic lesions on the lung surfaces were measured and compared, and supports the gross observation that the KPF1-Antx hMAb treated group showed lower lesion values (13.85 to 31.51 %) than those of the Isotype-Antx control hMAb group (32.81 to 37.05 %) (Figure 4F).

**Figure 4.**
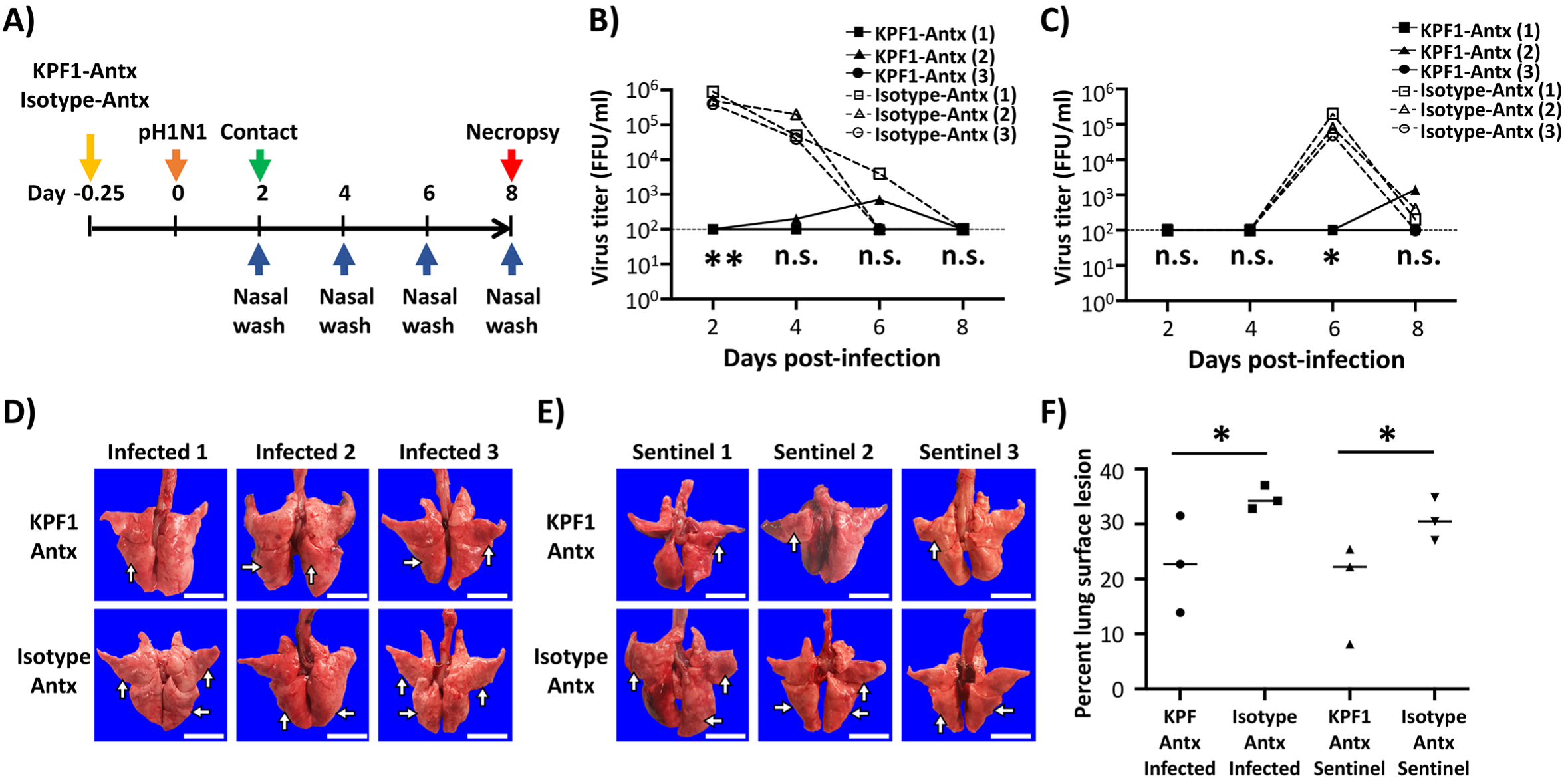
Prophylactic activity of KPF1-Antx hMAb in guinea pigs against pH1N1. **A) Schematic representation of the experimental approach:** Female Hartley guinea pigs (N = 3) were treated (i.p.) with 20 mg/kg of KPF1-Antx hMAb, or with 20 mg/kg of an IgG isotype control (Isotype-Antx). Six h post-treatment, guinea pigs were infected (i.n.) with 10^3^ PFU of pH1N1 and monitored daily for 8 days. At 1 d p.i., sentinel guinea pigs were introduced into the same cage of infected guinea pigs, allowing direct contact between the animals. **B-C) Viral titers from nasal washes:** Viral titers in the nasal washes of infected (**B**) and sentinel (**C**) guinea pigs were determined at 2, 4, 6, and 8 d p.i. by IFA (FFU/ml). The dotted line represents the limit of detection of the assay. * *p* < 0.05, ** *p* < 0.01, or no significance (n.s.). **D-E) Gross observation of lung pathology:** All animals were euthanized at 8 d p.i. and lungs were collected from infected (**D**) or sentinel (**E**) guinea pigs to observe gross pathological changes such as congestion and atelectasis (arrows). Scale bars = 1 cm. **F) Macroscopic pathology scoring:** Distributions of pathologic lesion such as consolidation, congestion, and atelectasis were measured using ImageJ and represented as the percent of the total lung surface area (%). * *p* < 0.05, or no significance (n.s.).

### 3.6. Therapeutic activity of KPF1-Antx hMAb in vivo

To assess the therapeutic activity of the KPF1-Antx hMAb, guinea pigs (N=3/group) were infected with 10^3^ PFU of pH1N1 and then treated, 24 h p.i., with 20 mg/kg of either KPF1-Antx or isotype-Antx control hMAbs. Sentinel guinea pigs (N=3/group) were put in contact with the infected guinea pigs at 2 d p.i. (Figure 5A) and evaluated for viral infection and shedding by determining viral titers in the nasal washes (Figures 5B and 5C, respectively). We observed high viral titers in infected guinea pigs treated with KPF1-Antx and the isotype-Antx control hMAbs at 2 d p.i. However, from 3 d p.i., the KPF1-Antx hMAb treated group showed lower levels of viral replication in one of the infected guinea pigs and no viral shedding in the two other infected guinea pigs as compared with the isotype-Antx-treated control group (Figure 5B). Importantly, no virus was detected in the sentinel guinea pig group that were put in contact with the infected KPF1-Antx hMAb (Figure 5C). As expected, gross pathologic observation of the KPF1-Antx hMAb treated group showed only mild locally extensive congestion and atelectasis in caudal lobes and surface lesion score ranging from 12.13 to 25.07 % (Figure 5F), while guinea pigs in the isotype-Antx hMAb treated group had more severe and larger distribution of pathologic lesions affecting middle and caudal lobes and surface lesion score, ranging from 34.03 to 37.20 %, (Figure 5F). Altogether, these results demonstrate that the plant-produced KPF1-Antx hMAb has both prophylactic and therapeutic activity against pH1N1, including the ability to prevent viral transmission, in guinea pigs.

**Figure 5.**
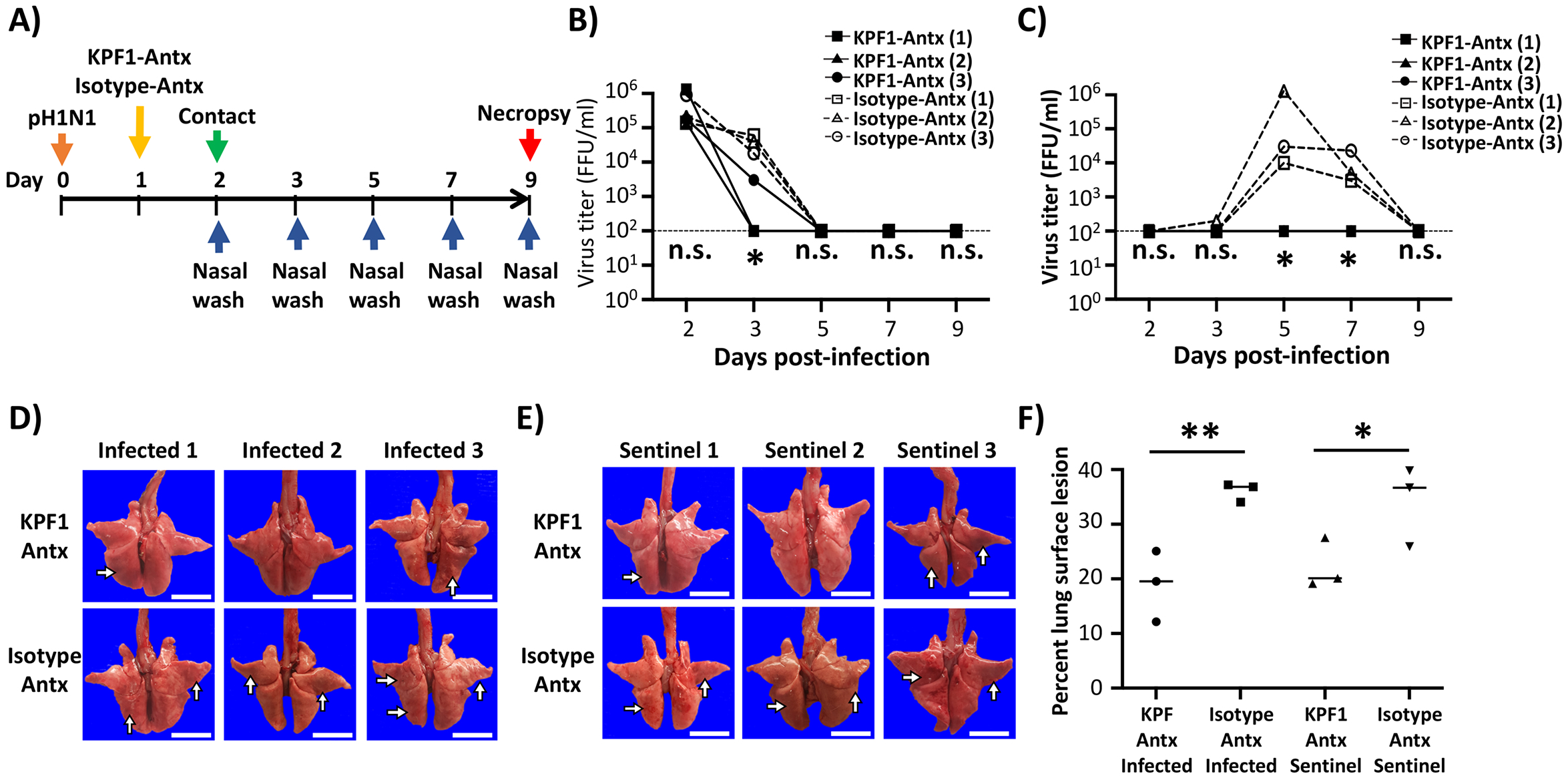
Therapeutic activity of KPF1-Antx hMAb in guinea pigs against pH1N1. **A) Schematic representation of the experimental approach:** Female Hartley guinea pigs (N = 3) were infected (i.n.) with 10^3^ PFU of pH1N1 and treated (i.p.) with 20 mg/kg of KPF1-Antx hMAb, or with 20 mg/kg of an IgG isotype control (Isotype-Antx) 24 h after infection. At 2 d p.i., sentinel guinea pigs (N = 3) were introduced into the same cages that infected guinea pigs, allowing direct contact between the animals. **B-C) Viral titers in nasal washes:** To measure viral titers in the upper respiratory tract, nasal washes of infected (**B**) and sentinel (**C**) animals were collected on 2, 3, 5, 7, and 9 d p.i. Viral titers were determined by IFA (FFU/ml). * *p* < 0.05, or no significance (n.s.). **D-E) Gross observation of lung pathology:** All animals were euthanized at 8 d p.i. and lungs were collected from infected (**D**) or sentinel (**E**) guinea pigs to observe gross pathological changes such as congestion and atelectasis (arrows). Scale bars = 1 cm. **F) Macroscopic pathology scoring:** Distributions of pathologic lesion, including consolidation, congestion, and atelectasis were measured using ImageJ and represented as the percent of the total lung surface area (%).* *p* < 0.05, ** *p* < 0.01, or no significance (n.s.).

## 4. Discussion

To date, FDA-licensed vaccines and antivirals are the most effective methods to prevent and/or control influenza viral infections. However, influenza vaccines require at least 2 weeks after administration to induce protective immune responses against viral infection. In the case of antivirals, their effectiveness is dependent on being administered within 48 h after appearance of symptoms [59, 60]. Importantly, antiviral-resistant strains have been described [1, 24–26]. Consequently, these prevention and treatment methods still leave substantial public health vulnerabilities.

In a previous study, we described a HEK293T-produced KPF1 hMAb (KPF1-HEK) that shows broad binding and neutralizing activity against H1N1 IAV isolates *in vitro* and potent prophylactic and therapeutic activities *in vivo* [19], because of its ability to bind to a conserved residue in the H1 hemagglutinin globular head of IAV [19]. In our current study, we examine the efficacy of a plant-produced KPF1 hMAb (KPF1-Antx) that possesses similar *in vitro* and *in vivo* properties to KPF1-HEK hMAb. KPF1-Antx was produced in *N. benthamiana* plants using the *vivo*XPRESS^®^ platform (Figure 1A), where four-week old plants were infiltrated with Agrobacteria containing expression vectors. Purified hMAb was recovered just seven days after infiltration from the foliage using standard antibody purification techniques. SDS-PAGE (Figure 2A) and SEC-HPLC (Figure 1C) revealed that KPF1-Antx and KPF1-HEK were highly similar. KPF1-Antx was engineered to have predominantly one glycan species – GnGn (Figure 1D). KPF1-Antx hMAb bound various recombinant and native H1 HAs as determined by ELISA and IFA assays (Figure 2 and 3, respectively). Importantly, KPF1-Antx hMAb showed neutralization and HAI activities similar to those of KPF1-HEK hMAb (Table1), highlighting that KPF1-Antx hMAb possesses and maintains similar avidity and affinity properties as the mammalian-produced KPF1 hMAb. Notably, judging by the potent binding affinity for recent isolates, such as ChCh/H1N1 and St. Petersburg27/H1N1 (Figure 2), KPF1 hMAb covers substantial antiviral breadth, including recent H1 isolates (Figure 3).

We also examined the activity of KPF1-Antx hMAb to protect (prophylactic) and treat (therapeutic) pH1N1 infected guinea pigs, as well as to prevent direct contact transmission (Figures 4 and 5, respectively). KPF1-Antx hMAb showed potent prophylactic and therapeutic activity against pH1N1 infection in guinea pigs (Figures 4 and 5, respectively), consistent with its *in vitro* antiviral and HAI properties. Moreover, administration of KPF1-Antx hMAb blocked transmission of pH1N1 infection to sentinel guinea pigs from direct contact with infected animals (Figures 4 and 5, respectively).

Previous studies described tobacco-derived plantibodies that showed potent therapeutic activity against multiple viruses such as EBOV, HBV, HIV, PEDV, RABV, WNV, and others in small animal models of infection, e.g. mice [40, 43–47]. However, to our knowledge, this is the first study demonstrating the ability of plantibodies to protect against influenza viral infections. Concerns exist that plantibodies can be immunogenic and/or allergenic in animals and humans because of a non-mammalian glycosylation pattern [61]. In view of that, a proprietary *N. benthamiana* plant line, KDFX, with knock-down of β1,2-xylosyltransferase and α1,3-fucosyltransfere, was used to reduce the addition of plant-specific glycan species. As a result, no plant-specific sugars were detected (Figure 1D). In the future, the *vivo*XPRESS^®^ platform could be used to tailor the N-glycan profile (i.e. addition of galactose and/or α1,6-fucose). Notably, none of the hMAb KPF1-Antx-or isotype-Antx-treated guinea pigs showed any adverse side effects and/or clinical signs in any of the in vivo experiments at a dose of 20 mg/kg.

Altogether, we substantiate the clinical potential of KPF1 to prevent and treat H1N1 infection in a second animal model, the guinea pig, including its potent ability to prevent transmission and for the first time, demonstrate the feasibility of using a plant-produced hMAb for the prophylactic and therapeutic treatment of influenza infections. Plant-produced hMAbs could represent an excellent option of the treatment and control of influenza viruses against which vaccines are not effective (e.g. seasonal) or available (e.g. pandemic) or FDA-approved antivirals are ineffective. Moreover, our results also suggest the feasibility of implementing hMAbs produced in tobacco plants for the treatment of other viral infections. However, further investigation will be needed to properly evaluate the safety and efficacy of using such plant-produced hMAbs in humans.

## Acknowledgments and funding

We thank the Biodefense and Emerging Infectious Research Resources Repository (BEI Resources) for providing reagents. This research was partially funded by the New York Influenza Center of Excellence (NYICE), a member of the National Institute of Allergy and Infectious Diseases (NIAID), National Institutes of Health (NIH), Department of Health and Human Services, Centers of Excellence for Influenza Research and Surveillance (CEIRS) contract No. HHSN272201400005C (NYICE), by the NIH/NIAID 1R01AI145332, and by the University of Rochester Technology Development funds to LM-S and JK.

## Author Contributions

LM-S and JK conceived and designed the experiments; J-GP, CY, MSP and AN performed the experiments and analyzed the data; HW, MS, AM produced and characterized KPF1-Antx; J-GP and CY wrote the original draft; all the authors reviewed and edited the paper.

## Conflicts of Interest

The authors declare no conflict of interests.

## References

1. Knipe, D. M.; Howley, P. M.; Cohen, J. I.; Griffin, D. E.; Lamb, R. A.; Martin, M. A.; Racaniello, V. R.; Roizman, B., Fields Virology. 6th editioin ed.; Wolters Kluwer Health: Philadelphia PA., 2013.

2. Li, K. S.; Guan, Y.; Wang, J.; Smith, G. J.; Xu, K. M.; Duan, L.; Rahardjo, A. P.; Puthavathana, P.; Buranathai, C.; Nguyen, T. D.; Estoepangestie, A. T.; Chaisingh, A.; Auewarakul, P.; Long, H. T.; Hanh, N. T.; Webby, R. J.; Poon, L. L.; Chen, H.; Shortridge, K. F.; Yuen, K. Y.; Webster, R. G.; Peiris, J. S., Genesis of a highly pathogenic and potentially pandemic H5N1 influenza virus in eastern Asia. Nature 2004, 430, (6996), 209–13.

3. Barr, I. G.; McCauley, J.; Cox, N.; Daniels, R.; Engelhardt, O. G.; Fukuda, K.; Grohmann, G.; Hay, A.; Kelso, A.; Klimov, A.; Odagiri, T.; Smith, D.; Russell, C.; Tashiro, M.; Webby, R.; Wood, J.; Ye, Z.; Zhang, W.; Writing Committee of the World Health Organization Consultation on Northern Hemisphere Influenza Vaccine Composition, f., Epidemiological, antigenic and genetic characteristics of seasonal influenza A(H1N1), A(H3N2) and B influenza viruses: basis for the WHO recommendation on the composition of influenza vaccines for use in the 2009-2010 northern hemisphere season. Vaccine 2010, 28, (5), 1156–67.

4. Clark, A. M.; DeDiego, M. L.; Anderson, C. S.; Wang, J.; Yang, H.; Nogales, A.; Martinez-Sobrido, L.; Zand, M. S.; Sangster, M. Y.; Topham, D. J., Antigenicity of the 2015–2016 seasonal H1N1 human influenza virus HA and NA proteins. PloS one 2017, 12, (11), e0188267.

5. Federici, C.; Cavazza, M.; Costa, F.; Jommi, C., Health care costs of influenza-related episodes in high income countries: A systematic review. PLoS One 2018, 13, (9), e0202787.

6. Molinari, N. A.; Ortega-Sanchez, I. R.; Messonnier, M. L.; Thompson, W. W.; Wortley, P. M.; Weintraub, E.; Bridges, C. B., The annual impact of seasonal influenza in the US: measuring disease burden and costs. Vaccine 2007, 25, (27), 5086–5096.

7. Neumann, G.; Kawaoka, Y., Transmission of influenza A viruses. Virology 2015, 479–480, 234-46.

8. Herfst, S.; Imai, M.; Kawaoka, Y.; Fouchier, R. A., Avian influenza virus transmission to mammals. Curr Top Microbiol Immunol 2014, 385, 137–55.

9. De Clercq, E., Antiviral agents active against influenza A viruses. Nat Rev Drug Discov 2006, 5, (12), 1015–25.

10. Mifsud, E. J.; Hayden, F. G.; Hurt, A. C., Antivirals targeting the polymerase complex of influenza viruses. Antiviral Res 2019, 169, 104545.

11. O’Hanlon, R.; Shaw, M. L., Baloxavir marboxil: the new influenza drug on the market. Curr Opin Virol 2019, 35, 14–18.

12. Su, S.; Fu, X.; Li, G.; Kerlin, F.; Veit, M., Novel Influenza D virus: Epidemiology, pathology, evolution and biological characteristics. Virulence 2017, 8, (8), 1580–1591.

13. Baker, S. F.; Nogales, A.; Finch, C.; Tuffy, K. M.; Domm, W.; Perez, D. R.; Topham, D. J.; Martinez-Sobrido, L., Influenza A and B virus intertypic reassortment through compatible viral packaging signals. J Virol 2014, 88, (18), 10778–91.

14. Influenza Research Database. https://www.fludb.org/brc/home.spg?decorator=influenza (Decemher 23th, 2019),

15. Parrish, C. R.; Kawaoka, Y., The origins of new pandemic viruses: the acquisition of new host ranges by canine parvovirus and influenza A viruses. Annu Rev Microbiol 2005, 59, 553–86.

16. Parrish, C. R.; Murcia, P. R.; Holmes, E. C., Influenza virus reservoirs and intermediate hosts: dogs, horses, and new possibilities for influenza virus exposure of humans. J Virol 2015, 89, (6), 2990–4.

17. Tong, S.; Zhu, X.; Li, Y.; Shi, M.; Zhang, J.; Bourgeois, M.; Yang, H.; Chen, X.; Recuenco, S.; Gomez, J.; Chen, L. M.; Johnson, A.; Tao, Y.; Dreyfus, C.; Yu, W.; McBride, R.; Carney, P. J.; Gilbert, A. T.; Chang, J.; Guo, Z.; Davis, C. T.; Paulson, J. C.; Stevens, J.; Rupprecht, C. E.; Holmes, E. C.; Wilson, I. A.; Donis, R. O., New world bats harbor diverse influenza A viruses. PLoS Pathog 2013, 9, (10), e1003657.

18. Tong, S.; Li, Y.; Rivailler, P.; Conrardy, C.; Castillo, D. A.; Chen, L. M.; Recuenco, S.; Ellison, J. A.; Davis, C. T.; York, I. A.; Turmelle, A. S.; Moran, D.; Rogers, S.; Shi, M.; Tao, Y.; Weil, M. R.; Tang, K.; Rowe, L. A.; Sammons, S.; Xu, X.; Frace, M.; Lindblade, K. A.; Cox, N. J.; Anderson, L. J.; Rupprecht, C. E.; Donis, R. O., A distinct lineage of influenza A virus from bats. Proc Natl Acad Sci U S A 2012, 109, (11), 4269–74.

19. Nogales, A.; Piepenbrink, M. S.; Wang, J.; Ortega, S.; Basu, M.; Fucile, C. F.; Treanor, J. J.; Rosenberg, A. F.; Zand, M. S.; Keefer, M. C.; Martinez-Sobrido, L.; Kobie, J. J., A Highly Potent and Broadly Neutralizing H1 Influenza-Specific Human Monoclonal Antibody. Sci Rep 2018, 8, (1), 4374.

20. Blanco-Lobo, P.; Nogales, A.; Rodriguez, L.; Martinez-Sobrido, L., Novel Approaches for The Development of Live Attenuated Influenza Vaccines. Viruses 2019, 11, (2).

21. Martinez-Sobrido, L.; Peersen, O.; Nogales, A., Temperature Sensitive Mutations in Influenza A Viral Ribonucleoprotein Complex Responsible for the Attenuation of the Live Attenuated Influenza Vaccine. Viruses 2018, 10, (10).

22. Nogales, A.; Martinez-Sobrido, L., Reverse Genetics Approaches for the Development of Influenza Vaccines. Int J Mol Sci 2016, 18, (1).

23. Mifsud, E. J.; Hayden, F. G.; Hurt, A. C., Antivirals targeting the polymerase complex of influenza viruses. Antiviral Res 2019, (169), 104545.

24. Takashita, E.; Kawakami, C.; Morita, H.; Ogawa, R.; Fujisaki, S.; Shirakura, M.; Miura, H.; Nakamura, K.; Kishida, N.; Kuwahara, T.; Mitamura, K.; Abe, T.; Ichikawa, M.; Yamazaki, M.; Watanabe, S.; Odagiri, T.; On Behalf Of The Influenza Virus Surveillance Group Of, J., Detection of influenza A(H3N2) viruses exhibiting reduced susceptibility to the novel cap-dependent endonuclease inhibitor baloxavir in Japan, December 2018. Euro Surveill 2019, 24, (3).

25. Earhart, K. C.; Elsayed, N. M.; Saad, M. D.; Gubareva, L. V.; Nayel, A.; Deyde, V. M.; Abdelsattar, A.; Abdelghani, A. S.; Boynton, B. R.; Mansour, M. M.; Essmat, H. M.; Klimov, A.; Shuck-Lee, D.; Monteville, M. R.; Tjaden, J. A., Oseltamivir resistance mutation N294S in human influenza A(H5N1) virus in Egypt. J Infect Public Health 2009, 2, (2), 74–80.

26. Omoto, S.; Speranzini, V.; Hashimoto, T.; Noshi, T.; Yamaguchi, H.; Kawai, M.; Kawaguchi, K.; Uehara, T.; Shishido, T.; Naito, A.; Cusack, S., Characterization of influenza virus variants induced by treatment with the endonuclease inhibitor baloxavir marboxil. Sci Rep 2018, 8, (1), 9633.

27. He, G.; Massarella, J.; Ward, P., Clinical pharmacokinetics of the prodrug oseltamivir and its active metabolite Ro 64-0802. Clin Pharmacokinet 1999, 37, (6), 471–84.

28. Davies, B. E., Pharmacokinetics of oseltamivir: an oral antiviral for the treatment and prophylaxis of influenza in diverse populations. J Antimicrob Chemother 2010, 65 Suppl 2, ii5–ii10.

29. Horadam, V. W.; Sharp, J. G.; Smilack, J. D.; McAnalley, B. H.; Garriott, J. C.; Stephens, M. K.; Prati, R. C.; Brater, D. C., Pharmacokinetics of amantadine hydrochloride in subjects with normal and impaired renal function. Ann Intern Med 1981, 94, (4 pt 1), 454–8.

30. Ng, K. E., Xofluza (Baloxavir Marboxil) for the Treatment Of Acute Uncomplicated Influenza. P T 2019, 44, (1), 9–11.

31. Wycoff, K. L., Secretory IgA antibodies from plants. Curr Pharm Des 2005, 11, (19), 2429–37.

32. Rodriguez, M.; Perez, L.; Gavilondo, J. V.; Garrido, G.; Bequet-Romero, M.; Hernandez, I.; Huerta, V.; Cabrera, G.; Perez, M.; Ramos, O.; Leyva, R.; Leon, M.; Ramos, P. L.; Triguero, A.; Hernandez, A.; Sanchez, B.; Ayala, M.; Soto, J.; Gonzalez, E.; Mendoza, O.; Tiel, K.; Pujol, M., Comparative in vitro and experimental in vivo studies of the anti-epidermal growth factor receptor antibody nimotuzumab and its aglycosylated form produced in transgenic tobacco plants. Plant Biotechnol J 2013, 11, (1), 53–65.

33. Hiatt, A.; Cafferkey, R.; Bowdish, K., Production of antibodies in transgenic plants. Nature 1989, 342, (6245), 76–8.

34. Lai, H.; He, J.; Engle, M.; Diamond, M. S.; Chen, Q., Robust production of virus-like particles and monoclonal antibodies with geminiviral replicon vectors in lettuce. Plant Biotechnol J 2012, 10, (1), 95–104.

35. Yuan, Q.; Hu, W.; Pestka, J. J.; He, S. Y.; Hart, L. P., Expression of a functional antizearalenone single-chain Fv antibody in transgenic Arabidopsis plants. Appl Environ Microbiol 2000, 66, (8), 3499–505.

36. Choi, Y. S.; Moon, J. H.; Kim, T. G.; Lee, J. Y., Potent In Vitro and In Vivo Activity of Plantibody Specific for Porphyromonas gingivalis FimA. Clin Vaccine Immunol 2016, 23, (4), 346–52.

37. Nakanishi, K.; Narimatsu, S.; Ichikawa, S.; Tobisawa, Y.; Kurohane, K.; Niwa, Y.; Kobayashi, H.; Imai, Y., Production of hybrid-IgG/IgA plantibodies with neutralizing activity against Shiga toxin 1. PLoS One 2013, 8, (11), e80712.

38. Chaisri, U.; Chaicumpa, W., Evolution of Therapeutic Antibodies, Influenza Virus Biology, Influenza, and Influenza Immunotherapy. Biomed Res Int 2018, 2018, 9747549.

39. Sully, E. K.; Whaley, K. J.; Bohorova, N.; Bohorov, O.; Goodman, C.; Kim, D. H.; Pauly, M. H.; Velasco, J.; Hiatt, E.; Morton, J.; Swope, K.; Roy, C. J.; Zeitlin, L.; Mantis, N. J., Chimeric plantibody passively protects mice against aerosolized ricin challenge. Clin Vaccine Immunol 2014, 21, (5), 777–82.

40. Shafaghi, M.; Maktoobian, S.; Rasouli, R.; Howaizi, N.; Ofoghi, H.; Ehsani, P., Transient Expression of Biologically Active Anti-rabies Virus Monoclonal Antibody in Tobacco Leaves. Iran J Biotechnol 2018, 16, (1), e1774.

41. Kopertekh, L.; Meyer, T.; Freyer, C.; Hust, M., Transient plant production of Salmonella Typhimurium diagnostic antibodies. Biotechnol Rep (Amst) 2019, 21, e00314.

42. Nakanishi, K.; Morikane, S.; Ichikawa, S.; Kurohane, K.; Niwa, Y.; Akimoto, Y.; Matsubara, S.; Kawakami, H.; Kobayashi, H.; Imai, Y., Protection of Human Colon Cells from Shiga Toxin by Plant-based Recombinant Secretory IgA. Sci Rep 2017, 7, 45843.

43. Lai, H.; Engle, M.; Fuchs, A.; Keller, T.; Johnson, S.; Gorlatov, S.; Diamond, M. S.; Chen, Q., Monoclonal antibody produced in plants efficiently treats West Nile virus infection in mice. Proc Natl Acad Sci U S A 2010, 107, (6), 2419–24.

44. Rattanapisit, K.; Srijangwad, A.; Chuanasa, T.; Sukrong, S.; Tantituvanont, A.; Mason, H. S.; Nilubol, D.; Phoolcharoen, W., Rapid Transient Production of a Monoclonal Antibody Neutralizing the Porcine Epidemic Diarrhea Virus (PEDV) in Nicotiana benthamiana and Lactuca sativa. Planta Med 2017, 83, (18), 1412–1419.

45. Hernandez-Velazquez, A.; Lopez-Quesada, A.; Ceballo-Camara, Y.; Cabrera-Herrera, G.; Tiel-Gonzalez, K.; Mirabal-Ortega, L.; Perez-Martinez, M.; Perez-Castillo, R.; Rosabal-Ayan, Y.; Ramos-Gonzalez, O.; Enriquez-Obregon, G.; Depicker, A.; Pujol-Ferrer, M., Tobacco seeds as efficient production platform for a biologically active anti-HBsAg monoclonal antibody. Transgenic Res 2015, 24, (5), 897–909.

46. Dubey, K. K.; Luke, G. A.; Knox, C.; Kumar, P.; Pletschke, B. I.; Singh, P. K.; Shukla, P., Vaccine and antibody production in plants: developments and computational tools. Brief Funct Genomics 2018, 17, (5), 295–307.

47. Olinger, G. G., Jr.; Pettitt, J.; Kim, D.; Working, C.; Bohorov, O.; Bratcher, B.; Hiatt, E.; Hume, S. D.; Johnson, A. K.; Morton, J.; Pauly, M.; Whaley, K. J.; Lear, C. M.; Biggins, J. E.; Scully, C.; Hensley, L.; Zeitlin, L., Delayed treatment of Ebola virus infection with plant-derived monoclonal antibodies provides protection in rhesus macaques. Proc Natl Acad Sci U S A 2012, 109, (44), 18030–5.

48. Nogales, A.; Avila-Perez, G.; Rangel-Moreno, J.; Chiem, K.; DeDiego, M. L.; Martinez-Sobrido, L., A novel fluorescent and bioluminescent Bi-Reporter influenza A virus (BIRFLU) to evaluate viral infections. J Virol 2019.

49. Nogales, A.; Baker, S. F.; Martinez-Sobrido, L., Replication-competent influenza A viruses expressing a red fluorescent protein. Virology 2015, 476, 206–16.

50. Nogales, A.; Rodriguez, L.; DeDiego, M. L.; Topham, D. J.; Martinez-Sobrido, L., Interplay of PA-X and NS1 Proteins in Replication and Pathogenesis of a Temperature-Sensitive 2009 Pandemic H1N1 Influenza A Virus. J Virol 2017, 91, (17).

51. Wing-Fai Cheung, H. W., Rebecca Pastora, Hirak Saxena, Warren W.; Wakarchuk, D. C. a. M. D. M., Development of fine-control expression vectors for post-translational modification of therapeutic proteins in plants. Molecular Biotechnology 2020, Submitted.

52. Castilho, A.; Beihammer, G.; Pfeiffer, C.; Goritzer, K.; Montero-Morales, L.; Vavra, U.; Maresch, D.; Grunwald-Gruber, C.; Altmann, F.; Steinkellner, H.; Strasser, R., An oligosaccharyltransferase from Leishmania major increases the N-glycan occupancy on recombinant glycoproteins produced in Nicotiana benthamiana. Plant Biotechnol J 2018, 16, (10), 1700–1709.

53. Garabagi, F.; McLean, M. D.; Hall, J. C., Transient and stable expression of antibodies in Nicotiana species. Methods Mol Biol 2012, 907, 389–408.

54. Jensen, E. C., Quantitative analysis of histological staining and fluorescence using ImageJ. Anat Rec (Hoboken) 2013, 296, (3), 378–81.

55. National Research Council (U.S.). Committee for the Update of the Guide for the Care and Use of Laboratory Animals, I. f. L. A. R. U. S., Guide for the Care and Use of Laboratory Animals. In edn, t., Ed. National Academies Press: Washington (DC), 2011.

56. Gauger, P. C.; Loving, C. L.; Khurana, S.; Lorusso, A.; Perez, D. R.; Kehrli, M. E., Jr.; Roth, J. A.; Golding, H.; Vincent, A. L., Live attenuated influenza A virus vaccine protects against A(H1N1)pdm09 heterologous challenge without vaccine associated enhanced respiratory disease. Virology 2014, 471–473, 93-104.

57. Morgan, S. B.; Hemmink, J. D.; Porter, E.; Harley, R.; Shelton, H.; Aramouni, M.; Everett, H. E.; Brookes, S. M.; Bailey, M.; Townsend, A. M.; Charleston, B.; Tchilian, E., Aerosol Delivery of a Candidate Universal Influenza Vaccine Reduces Viral Load in Pigs Challenged with Pandemic H1N1 Virus. J Immunol 2016, 196, (12), 5014–23.

58. Robert, S.; Goulet, M. C.; D’Aoust, M. A.; Sainsbury, F.; Michaud, D., Leaf proteome rebalancing in Nicotiana benthamiana for upstream enrichment of a transiently expressed recombinant protein. Plant Biotechnol J 2015, 13, (8), 1169–79.

59. Nicholson, K. G.; Aoki, F. Y.; Osterhaus, A. D.; Trottier, S.; Carewicz, O.; Mercier, C. H.; Rode, A.; Kinnersley, N.; Ward, P., Efficacy and safety of oseltamivir in treatment of acute influenza: a randomised controlled trial. Neuraminidase Inhibitor Flu Treatment Investigator Group. Lancet 2000, 355, (9218), 1845–50.

60. McLean, H. Q.; Belongia, E. A.; Kieke, B. A.; Meece, J. K.; Fry, A. M., Impact of Late Oseltamivir Treatment on Influenza Symptoms in the Outpatient Setting: Results of a Randomized Trial. Open Forum Infect Dis 2015, 2, (3), ofv100.

61. Gomord, V.; Fitchette, A. C.; Menu-Bouaouiche, L.; Saint-Jore-Dupas, C.; Plasson, C.; Michaud, D.; Faye, L., Plant-specific glycosylation patterns in the context of therapeutic protein production. Plant Biotechnol J 2010, 8, (5), 564–87.

